# Piezo1 Mutant Zebrafish as a Model of Idiopathic Scoliosis

**DOI:** 10.1101/2023.10.23.563500

**Authors:** Ramli, Toshihiro Aramaki, Masakatsu Watanabe, Shigeru Kondo

**Affiliations:** Laboratory of Pattern Formation, Graduate School of Frontier Biosciences, Osaka University, Japan

**Keywords:** idiopathic scoliosis, piezo1, TMD, vertebral bone, zebrafish

## Abstract

Scoliosis is a condition where the spine curves sideways, unique to humans due to their upright posture. However, the cause of this disease is not well understood because it’s challenging to find a model for experimentation. This study aimed to create a model for human idiopathic scoliosis by manipulating the function of mechanosensitive channels called Piezo channels in zebrafish. Zebrafish were chosen because they experience similar biomechanical forces to humans, particularly in relation to the role of mechanical force in scoliosis progression. Here we describe *piezo1* and *piezo2a* are involved in bone formation, with a double knockout resulting in congenital systemic malformations. However, an in-frame mutation of *piezo1* led to fully penetrant juvenile-onset scoliosis, bone asymmetry, reduced tissue mineral density, and abnormal intervertebral discs-resembling non-congenital scoliosis symptoms in humans. These findings suggest that functional Piezo channels responding to mechanical forces are crucial for bone formation and maintaining spine integrity, providing insights into skeletal disorders.

## 1. Introduction

Scoliosis is a spinal deformity in which the spine curves sideways. It is estimated that approximately 5% of the world population is affected by scoliosis, the most common being idiopathic scoliosis that develops during adolescence (Konieczny et al., 2013). Various studies have linked the incidence of idiopathic scoliosis to central nervous system abnormalities (Burwell et al., 2009), hormonal imbalances (Kulis et al., 2015), muscle defects (Allam and Schwabe, 2013), and genetics (Wise et al., 2008), but details regarding the mechanisms of pathogenesis and other factors are not yet known.

One factor that has received attention as a possible cause of spinal deformity is mechanical forces (pressure and shear stress) (Hawes and O’brien, 2006). When the spine is subjected to unbalanced forces, the intervertebral discs are asymmetrically compressed and unequal growth of the spine occurs. This unequal spinal growth is thought to cause a positive feedback that results in a continual imbalance of mechanical loading, which ultimately inhibits spinal growth (Xie et al., 2022).

Animal models are commonly used to study pathology. However, in the case of scoliosis, the direction of gravity on the spine is different, making common animal models such as mice and rats unsuitable for research. In humans, due to the upright posture, the gravitational load is applied from a direction parallel to the spinal column (Castelein et al., 2005). In contrast, in quadrupedal animals, the direction of the vertebrae and the direction of gravity are perpendicular, so the contribution of gravity to bone growth is quite different (Xie et al., 2022). Therefore, bipedal chicks have been tried as an animal model for scoliosis, but they are not a perfect model for the condition because their spinal columns are not as flexible as those of humans, limiting spinal flexion and extension (Fagan et al., 2009).

Interestingly, on the other hand, guppies, a fish species, are known to spontaneously develop scoliosis. Fish are similar to humans in that during swimming, caudal propulsive forces cause the spine to compress in a direction similar to that of humans (Gorman and Breden, 2009). The fact that several features of scoliosis in guppies have been found to be similar to those of humans also suggests the possibility of using fish as a pathological model of scoliosis (Gorman et al., 2007). If fish could be used as a model for the development of scoliosis, zebrafish, which can be easily genetically modified, would be more experimentally convenient than guppies.

There are several genetic lineages in people presumed to be related to scoliosis, one of which is the mechanoreceptor Piezo channels. Piezo channels are located on the cell membrane and act as mechanotransducers. When physical forces are applied, Piezo channels are activated and cations such as calcium move into the cell, and activate several signaling pathways. There are two types of Piezo channels, Piezo1 and Piezo2 channels, which are encoded by the *piezo1* and *piezo2* genes, respectively. Piezo1 channel is highly expressed in non-sensory tissues such as the lung, skin, and bladder, unlike Piezo2, which is highly expressed in sensory tissues (Murthy et al., 2017).

It has been confirmed that *piezo1* genes are highly expressed in mesenchymal stem cells, osteoblasts, chondrocytes and nucleus pulposus cells of intervertebral disc of the spine (Zhu et al., 2021). This underlines that *piezo1* might contribute to several bone metabolic and degenerative diseases such as osteoporosis, idiopathic scoliosis, or intervertebral disc degeneration. In a recent report, exome analysis revealed novel mutations in PIEZO1 in patients with primary lymphatic dysplasia with symptoms of facial bone hypoplasia, thoracolumbar scoliosis, and short stature (Lee et al., 2021).

In this study, we proposed that optimal sensing of mechanical force is crucial for the pathogenesis of scoliosis, not just the force itself. To investigate this, we disrupted the ability of zebrafish to sense mechanical force by knocking out *piezo* genes. Zebrafish have three *piezo* genes in their genome (*piezo1, piezo2a*, and *piezo2b*), and when we deleted both *piezo1* and *piezo2a* simultaneously or introduced an in-frame mutation of *piezo1*, the fish exhibited symptoms similar to bone diseases, including scoliosis and intervertebral disc degeneration. These findings indicate that zebrafish with deleted *piezo* genes could serve as an animal model for studying scoliosis. Our results provide valuable insights into the role of Piezo channels in bone development and may shed light on potential causes of skeletal disorders.

## 2. Material and methods

### 2.1 Zebrafish husbandry

All experiments in this study were conducted in accordance with the prescribed guidelines and approved protocols for the handling and use of animals at Osaka University. Tübingen (Tü) strain was used as the wildtype zebrafish and all fish were maintained under standard conditions. Fish were kept under a photoperiod of 14 h light/10 h darkness at 28 °C in a fish breeding room. Embryos were obtained by crossing mature males and females at 28 °C. Embryos were collected and incubated in a thermostatic incubator at 28.5 °C. After hatching, larvae were transferred to the fish breeding room and fed paramecia until they grew to be able to eat artificial diets.

### 2.2 CRISPR/Cas9

For generating zebrafish mutant, CRISPR/Cas9-mediated method was utilized. The sgRNA target site for each gene was designed using crisprscan.org. The sequences of the oligonucleotide for sgRNA are listed in supplementary table S1. To Knock-out *piezo1* and *piezo2a* gene, exon 5 of *piezo1* with target sequence 5’ CCTCAGGGTGTGCTGTGGCTCCT 3’ and exon 4 of *piezo2a* with target sequence 5’ TGGCCACGCTCATCCGCCTCTGG 3’ were chosen. For sgRNA transcription, the template DNA was made by polymerase chain reaction using oligonucleotide containing target exon and tracrRNA backbone. Then, about 50 ng synthesized template DNA was *in vitro* transcribed using *in vitro* Transcription T7 Kit (for siRNA Synthesis) (Takara bio), treated with DNase I (Takara Bio), and purified by RNA clean and concentration kit (Zymo Research).

To generate *piezo1* in-frame mutant, an effective single guide RNA (sgRNA) sequence for zebrafish *piezo1* had been reported previously (Gudipaty et al., 2017). This sgRNA targeted the splice acceptor sequence of exon 2 of zebrafish *piezo1* gene. The target sequence is 5’-cagCCTGCATATTTCGCTAC-3’ (exonic sequence is shown in uppercase, intronic sequence in lowercase). The 20 nucleotides sequence upstream of PAM were cloned into pDR274 plasmid containing RNA loop structure sequence required for recognition by Cas9 enzyme, and T7 promoter sequence that allowed for *in vitro* synthesis of sgRNA using MEGA Script T7 Kit (Invitrogen #AM1334). Synthesized sgRNA was mixed with Cas9 protein (NEB# M0646T) just before microinjection into zebrafish embryo.

### 2.3 Microinjection and genotyping

One-cell stage zebrafish embryos were injected with 1-2 nl of injection solution containing 300 ng/μl of Cas9 enzyme (NEB# M0646T) and 12.5 ng//μl of sgRNA. For genotyping, the DNA was extracted from embryos at 3 dpf or amputated caudal fin at adult stage. These samples then were incubated in DNA extraction buffer supplemented with Proteinase K at 55 °C for 2 hours and 95 °C for 5 minutes. The extracted samples were diluted 300 times using distilled water and about 1 μl of diluted sample was used as a template for 20 μl standard PCR amplification. Primers for genotyping are listed in supplementary table S2.

### 2.4 Micro-CT

To analyze bone morphology and phenotypes in detail, utilize micro–computed tomography (micro-CT) is used. Whole-body fish was fixed in 4% PFA overnight. Fixed samples were observed by micro–CT□scanner SkyScan 1,172 (SkyScan NV, Aartselaar, Belgium) following the manufacturer’s instructions. The X□ray source ranged from 50□kV, and the datasets were acquired at a resolution of 5 μm/pixel for abdominal or caudal part and 12 □μm/pixel for whole body, depending on the size of each vertebral body. Samples were scanned in air dry condition. Neural arches and hemal arches were not included in bone analysis. After applying a fixed threshold for all samples, 3D evaluation is conducted using CTVox and CTAn. Orthoslice images were obtained using Dataviewer (Bruker, Kontich, Belgium).

### 2.5 Alizarin Red S staining

For live imaging, fish at larvae (8 dpf), juvenile (22 dpf), and young adult (35 dpf) were incubated in 0.005% alizarin red S (Sigma) overnight and washed three times with tank water. Before observation, samples were anesthetized using Ethyl 3-aminobenzoate methanesulfonate. Imaging was performed on a BZ-X710 Keyence fluorescence microscope.

### 2.6 RNA extraction and cDNA synthesis

To perform relative mRNA expression analysis, total RNA was extracted from vertebral bone using RNeasy Lipid tissue mini kit (QIAGEN) according to the manufacturer′s protocol. Subsequently, RNA concentration was measured using Q5000 micro-volume spectrophotometer (TOMY). About 500 ng of extracted RNA was used for cDNA synthesized using ReverTra Ace qPCR RT kit following the manufacturer′s protocol. The cDNA was stored at -20 °C until use.

### 2.7 Cell-attached patch clamp mode

To confirm whether the 11 amino acid deletion affects Piezo1 channel function, we performed patch-clamp experiment using cell-attached mode. The cDNAs of zebrafish wildtype-*piezo1* and 11-amino acids deletion mutant were cloned into pIRES2-GFP vector, respectively. The constructed plasmids then were purified using nucleospin plasmid-transfection grade (TAKARA) and transfected into piezo1-deficient N2A cells (Sugisawa et al., 2020). About 48 to 72 hours after transfection, the channel activities were recorded.

Briefly, the recordings were conducted at room temperature with a mean pipette resistance of 4.0 to 6.0 MΩ. In this recording, the pipette solution contained 140 mM NaCl, 5 mM KCl, 2 mM MgCl_2_, 2 mM CaCl_2_, 10 mM HEPES (pH 7.4 adjusted with NaOH), and 10 mM glucose same with bath solution. Following giga-seal formation, the holding potential was clamped to -80 mV and the channel was stimulated by applying a negative pressure of -40mmHg.

### 2.8 qPCR

The primers for this experiment are listed in supplementary table S3. To perform qPCR analysis, SYBR Green PCR Master Mix (Applied Biosystem) was used and carried out in the Applied Biosystems™ StepOne™ Real-Time PCR System. The amplification conditions were 95 °C for 10 min, 40 cycles at 95 °C for 15 s, and 60 °C for 1 min. All reactions were performed in triplicate. Relative mRNA expressions were normalized to *beta-actin* (*actb1*) gene. Primer sets for qPCR were listed in supplementary table S3.

### 2.9 Rescue experiment with introducing functional piezo1

Zebrafish *piezo1* coding sequence (CDS) was amplified by PCR and then cloned into the pTol2 plasmid with *osterix* promoter (Kawakami et al., 2000;Renn and Winkler, 2009). Sequences of the primers used to amplify *piezo1* CDS are piezo1-F1_Sal1; 5-TTTTGGCAAAGAATTGTCGACCACCATGGAGCTTCAGGTGGTA-3’ and piezo1-R1_Not1; 5’-CGTTAGGGGGGGGGGGCGGCCGCTCAGTTGTGATTCTTCTCTC-3’. In addition, since Piezo1 protein has many hydrophobic domains that cause toxicity in *E. coli*, intron 10 and intron 42 were inserted in CDS. Purified plasmid was mixed with *in vitro* synthesized Tol2 Transposase mRNA (10□ng/μL each), and the mixture was injected into fertilized eggs (1 nL/egg) at the single-cell stage.

### 2.10 Statistical analysis

All data are presented as mean ± SD. Statistical analysis was performed by two-tailed Student’s t-test for the two groups and by one-way analysis of variance (ANOVA) test and Tukey’s post-test for the three groups. Differences were considered statistically significant at *P < 0*.*05*.

## 3. Result

### 3.1 Generation of piezo null mutant

Zebrafish have three different *piezo* genes: *piezo1* (XM_691263), *piezo2a* (XM_021470255), and *piezo2b* (XM_021468270). In whole-mount *in situ* hybridization using 48hpf larval zebrafish, both expressions of *piezo1* and *piezo2a* were detected in many tissues, including bone, while *piezo2b* expression was limited to cells of the nervous system (Faucherre et al., 2013). This suggests that zebrafish *piezo1* and *piezo2a* may have redundant functions.

To generate *piezo* activity-deficient zebrafish, we utilized the CRISPR/Cas9 method (Ran et al., 2013). A guide RNA targeting the internal sequence of exon 5 of *piezo1* gene or exon 4 of *piezo2a* gene was synthesized and co-injected with Cas9 mRNA into one-cell stage wildtype embryos. Following the screening of several founders that transmitted to the F1 progeny, we established two independent stable lines of each mutant gene. We verified the sequence of each mutant allele and confirmed that each mutant gene generated a premature termination codon (PTC) as a result of an altered mRNA reading frame (Supplementary Figure S1).

### 3.2 Phenotypes of the zebrafish with double mutations of *piezo1* and *piezo2a*

As we expected from the similarity of expression pattern, the single mutants of *piezo1*^*-/-*^ and *piezo2a*^*-/-*^ were viable and fertile. Neither mutant showed morphological abnormalities in the larval and juvenile stages (Supplementary Figure S2).

Next, to obtain large numbers of fish with double mutations, fish sharing the *piezo1*^*-/+*^; *piezo2a*^*-/-*^ genotype were crossed. In this case, 2-30% of the individuals died around 10 days after fertilization. When the gene was examined, fish with the double homozygous mutation survived to day 8, but all died by day 11 (Supplementary Figure S3A). This indicates that the double mutant fish is lethal in the juvenile stage.

The surviving fish on day 8 showed a shortening of body length (Supplementary Figure S3B) and curvature of the body axis (Figure 1A,1B). In addition, the swim bladder was not fully inflated (90%) (Figure 1B arrowhead). Alizarin red S staining revealed that all double homozygous mutants showed inadequate bone formation such as small vertebral segment, hemivertebral and vertebral bone missing compared to their siblings (Figure 1C-1F). From these data, it is shown that Piezo channels are essential for bone formation and mineral deposition during early development. However, because of the lethality in the young stage, it is not suitable for pathological model of scoliosis.

**Figure 1.**
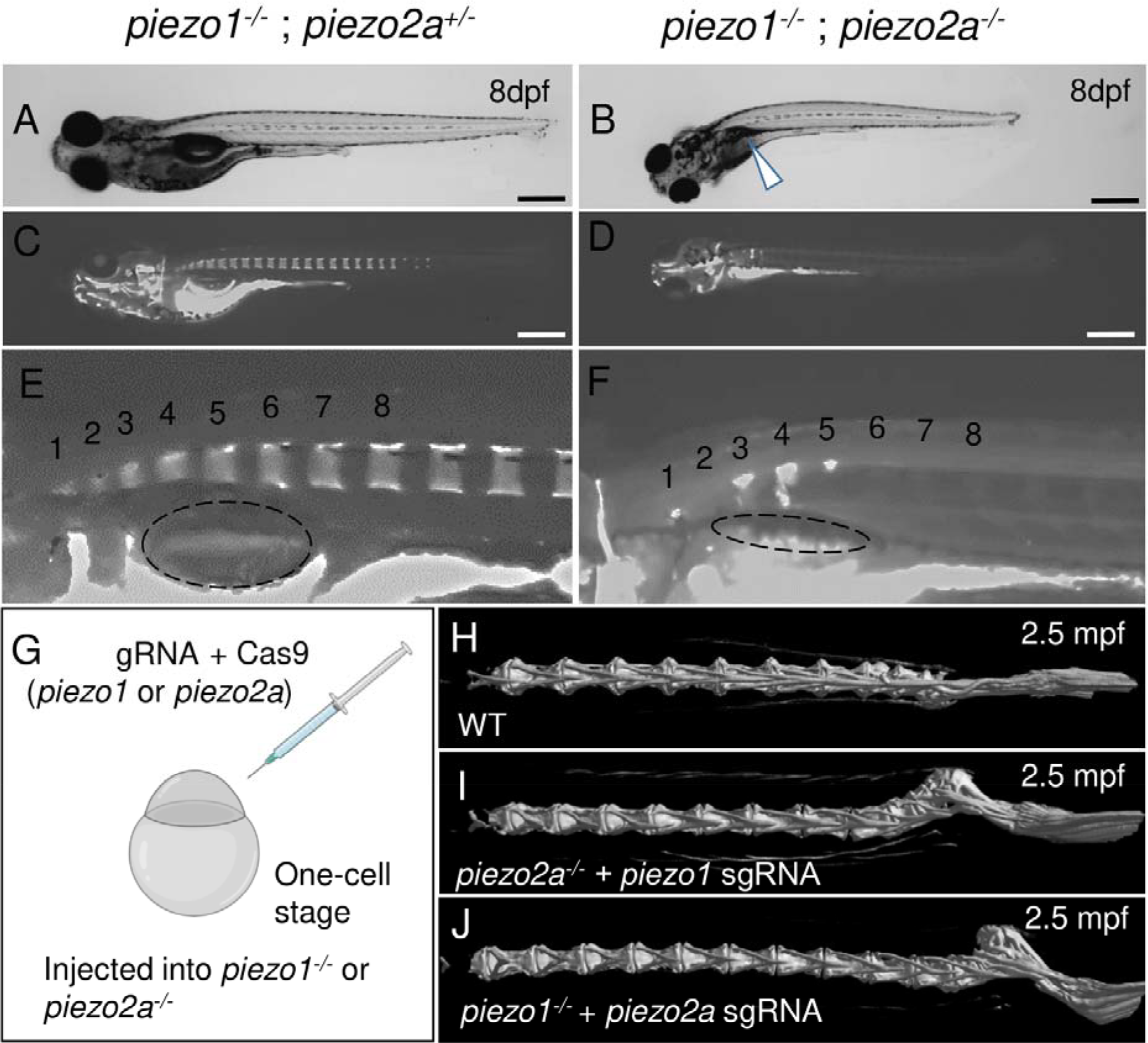
*piezo1* and *piezo2a* are essential for bone formation and mineral deposition in early stage of development. (A)(B) Gross phenotypes and (C)(D) mineralization patterns of sibling and *piezo1*^*-/-*^; *piezo2a*^*-/-*^ mutant at 8 days after fertilization (dpf). White arrow indicates uninflated swim bladder. Scale = 500 μm. (E)(F) Magnified images of the anterior body of sibling and double mutant. Numbers 1 to 8 indicate the position of each vertebral bone. The dashed circle indicates swim bladder. Notably, the swim bladder in double mutant was small and uninflated. (G) Schematic diagram of generating piezo transient KO mutant (CRISPRant). Comparison of 3D reconstruction of micro-CT images of vertebral bone at caudal part from (H) wildtype, (I) *piezo1* sgRNA injected to *piezo2a*^*-/-*^, and (J) *piezo2a* sgRNA injected to *piezo1*^*-/-*^ at 2.5 months after fertilization.

### 3.3 Mosaic mutant displays bone curvature

If the cause of death is an abnormality that occurs in an organ other than the spine, CRISPRants (F0 mutants), in which loss of function of the target gene occurs in a mosaic fashion, may avoid the lethality that occurs early in development and allow us to study effects during later developmental stages (She et al., 2019;Buglo et al., 2020). To test this possibility, we injected sgRNAs targeting *piezo1* or *piezo2a* into stage 1 cells in a *piezo2a* null mutant or *piezo1* null mutant background (Figure 1G).

As a result, no abnormal phenotypes were observed in either mutant background at the larval to juvenile stage. However, by day 17 post-fertilization, small curvature of the spine near the tail of the body occurred in more than 80% of *piezo2a*^*-/-*^ injected with *piezo1* sgRNA and about 50% of *piezo1*^*-/-*^ injected with *piezo2a* sgRNA (Figure 1H-1J; Supplementary Figure S4). This curvature was even more severe in adulthood, with the abnormal vertebrae located near the distal end of the tail and near the hypural complex.

In this mosaic mutants, the bone morphology mutation shares some similarities with scoliosis, particularly in its occurrence during the process of growth. This fact suggests that artificial mutations in the *piezo* genes may be able to create a pathological model of scoliosis, but in order to create a usable model, lethality must be avoided by simpler methods.

### 3.4 Generation of *piezo1* in-frame mutant

Next, we analyzed in-frame mutation of the *piezo1* gene. In the process of creating a deletion mutation in the *piezo1* gene, an in-frame mutation was obtained when a guide RNA targeting the splice acceptor sequence in exon 2 was used. In this allele, the removal of the splice acceptor site “AG” resulted in a deletion of 11 amino acids, without the insertion of a premature stop codon (Figure 2A,2B). The mutant was designated as *piezo1*^*11aa del/11aa del*^ and the resulting protein was named 11aa del-Piezo1. The 11 amino acid deletion region is conserved among vertebrates (Supplementary Figure 5) and likely plays an important role in Piezo1 channel function.

**Figure 2.**
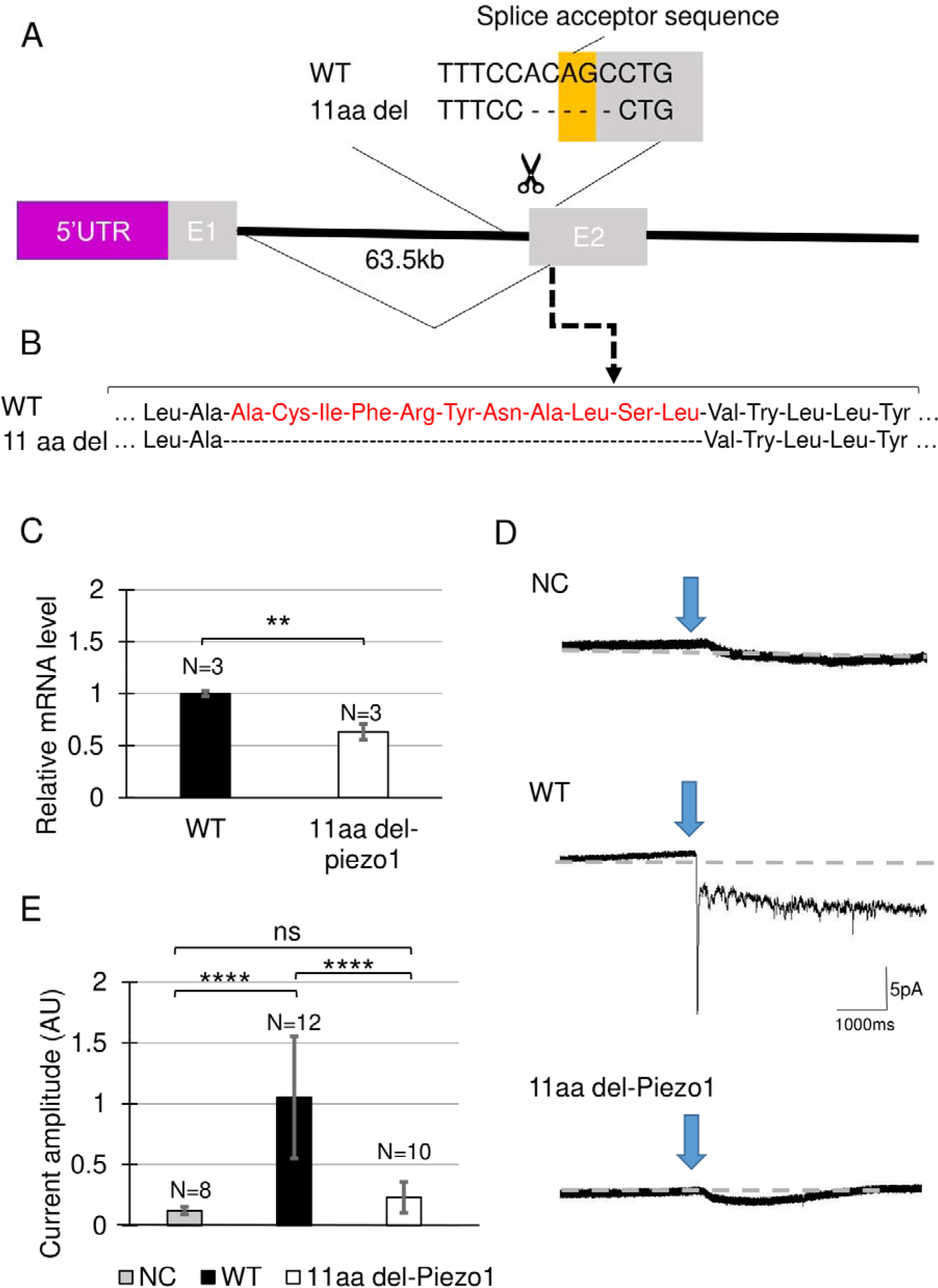
Generating *piezo1* in-frame mutant. (A) Schematic illustration of CRISPR target in *piezo1* gene with a guide RNA targeting splice acceptor site of exon2. Through this guideRNA, the splice acceptor site was completely eliminated, generating cryptic acceptor site which deleted several bases of internal exon 2. (B) Confirmation of amino acid sequence. (C) Quantification of relative mRNA level. Values are presented as mean ± SD and analyzed using student t-test. *\*\*P < 0*.*01*. (D) Representative traces of negative pressure-induced inward currents recorded at -80 mV in negative control (NC), zebrafish Piezo1 Wildtype (WT), and zebrafish 11aa del-Piezo1. Blue arrows indicate the time when negative pressure was applied. Grey dashed lines indicate baseline of the current. (E) Measurement of current amplitude, measured in arbitrary unit (AU). Value are presented as mean ±SD and analyzed using one-way ANOVA followed by Tukey’s test. *\*\*\*\* P < 0*.*0001*. ns indicates not significant.

First, we measured mRNA levels of 11aa del-Piezo1 using qPCR to determine whether *piezo1* gene in *piezo1*^*11aa del/11aa del*^ is transcribed normally. The results showed that the mRNA level of *piezo1* in *piezo1*^*11aa del/11aa del*^ mutant was reduced by almost half compared to the wildtype (Figure 2C).

Next, we assessed the channel activity of 11aa del-Piezo1 using a cell-attached patch clamp assay. The zebrafish *piezo1* genes were obtained from both the wildtype and mutant forms, inserted into an expression vector, and transfected into *piezo1*-deficient N2A cells. After about 48 hours of incubation, channel activity was recorded. It was found that in wildtype Piezo1, an inward current flowed when negative pressure was applied with a pipette, whereas only a very weak current flow was observed in the 11aa del-Piezo1 (Figure 2D, 2E). These results clearly indicated that the deletion of 11 amino acids in the N-terminal domain of Piezo1 caused a functional defect in the Piezo1 channel. In the following experiments, we examine the effects of this on spine formation in detail.

### 3.5 *piezo1* in-frame mutant (*piezo1*^*11aa del/11aa del*^) develops juvenile-onset scoliosis

At 4 days post-fertilization (dpf), there were no critical differences between wildtype and *piezo1*^*11aa del/11aa del*^. However, at later stages, during the larval and early juvenile period, the body length of the mutant fish became relatively shorter than that of wildtype, suggesting the late-onset abnormality in the vertebrae (Figure 3B).

**Figure 3.**
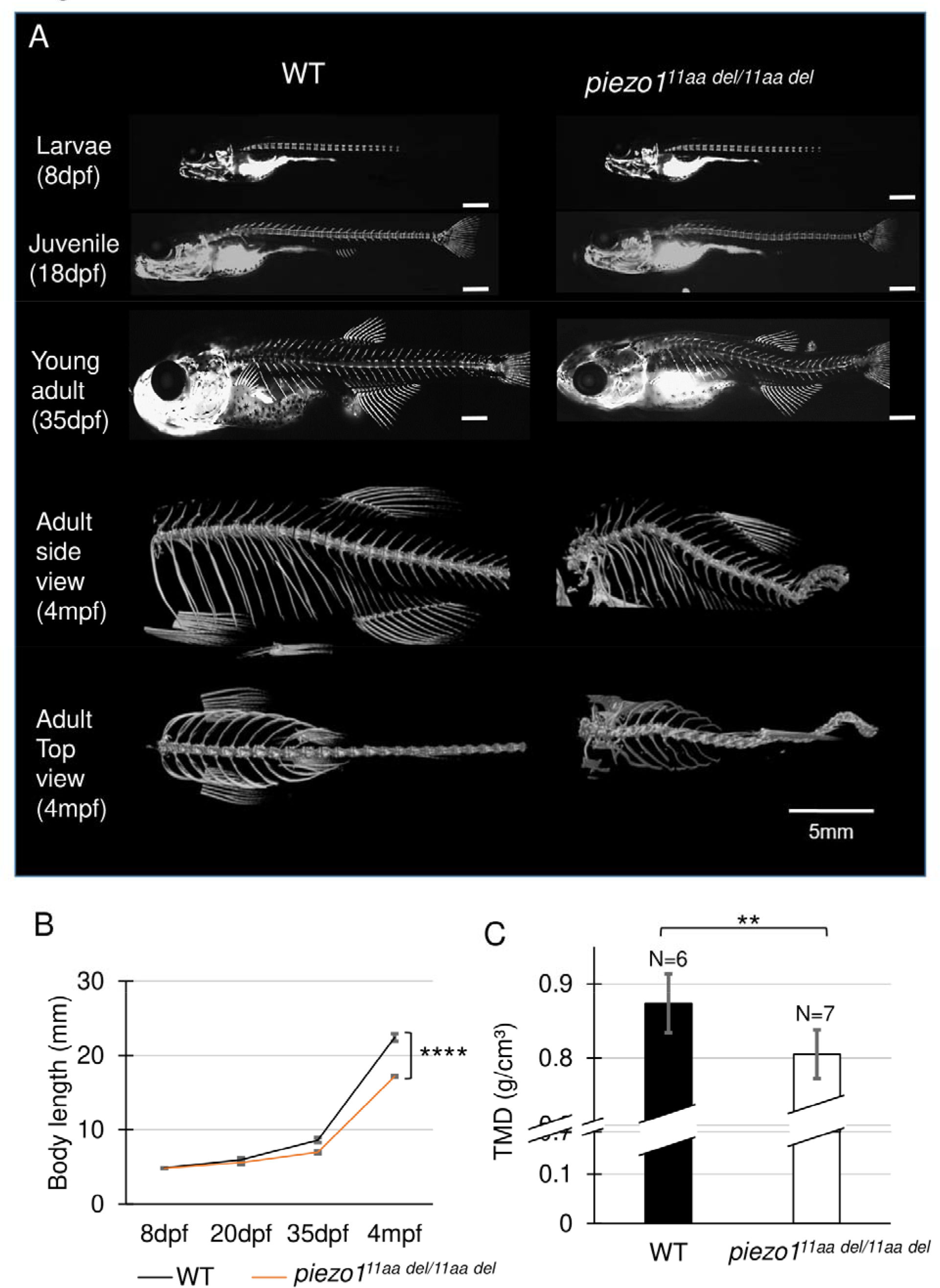
Phenotype of *piezo1* in-frame mutant (*piezo1*^*11aa del/11aa del*^). (A) Time course observation of mineralized bone using alizarin red S staining between wildtype and *piezo1*^*11aa del/11aa del*^ mutant from larvae (8 dpf) to young adult (35 dpf) (Used scale 500 μm) and 3D rendering of micro-CT images of wildtype and *piezo1*^*11aa del/11aa del*^ mutant at 4 months. (B) Graph depicting body length. (C) Measurement of Tissue Mineral Density (TMD). Value are presented as mean ± SD and analyzed using student t-test. *\*\* P < 0*.*01; **** P < 0*.*0001*.

Alizarin red staining to track vertebral formation at 8 dpf, 22 dpf, and 35 dpf showed that the spines gradually curved as the fish grew, and heterotopic bone formation was occurring in the intervertebral space. Unlike double mutant fish, however, there was no significant variation in the number or spacing of vertebrae or in the morphology of individual vertebrae (Figure 3A).

In adults, the curvature of the spine was even more severe, and it appears to be folded in particularly severe areas. This curvature occurs both laterally and vertically (Figure 3A). These symptoms resemble human scoliosis (Cheng et al., 2015).

We measured Tissue Mineral Density (TMD) from CT data to determine the degree of calcium deposition in bone. We found that *piezo1*^*11aa del/11aa del*^ at 3 months of age had predominantly lower TMD values than the wildtype (Figure 3C). This is consistent with various reports when they assessed the bone mineral density in scoliotic patients (Sarioglu et al., 2019;Li et al., 2020;Almomen et al., 2021).

### 3.6 Detailed observation of vertebra shape by CT

Next, micro-CT measurements were taken to examine the shape of the individual vertebrae and surrounding bone (Figure 4A,4B). As shown in Figure 3A, the vertebrae were generally curved, but there were no significant changes in the shape of the neural or haemal arches. To assess the deformity of individual vertebrae causing the curvature, the size of individual vertebrae was measured and expressed as the ratio of anterior/posterior height (H1/H2) and dorsal/ventral length (L1/L2) (Bearce et al., 2022). Figure 4 shows data for the 8th to 13th vertebrae of 5 individual fish. In the wildtype, both ratios were approximately 1, with very little variation among individuals or by vertebral position (Figure 4D,4E). In the mutant, however, there was variation for each individual vertebra, and the variation in length ratio (L1/L2) was considerably greater than the variation in height ratio (H1/H2) ((Figure 4F,4G). This fact suggests that the curvature of the spinal column has caused more longitudinal compression, which is also characteristic of scoliosis in humans (Shea et al., 2004;Schlager et al., 2018).

**Figure 4.**
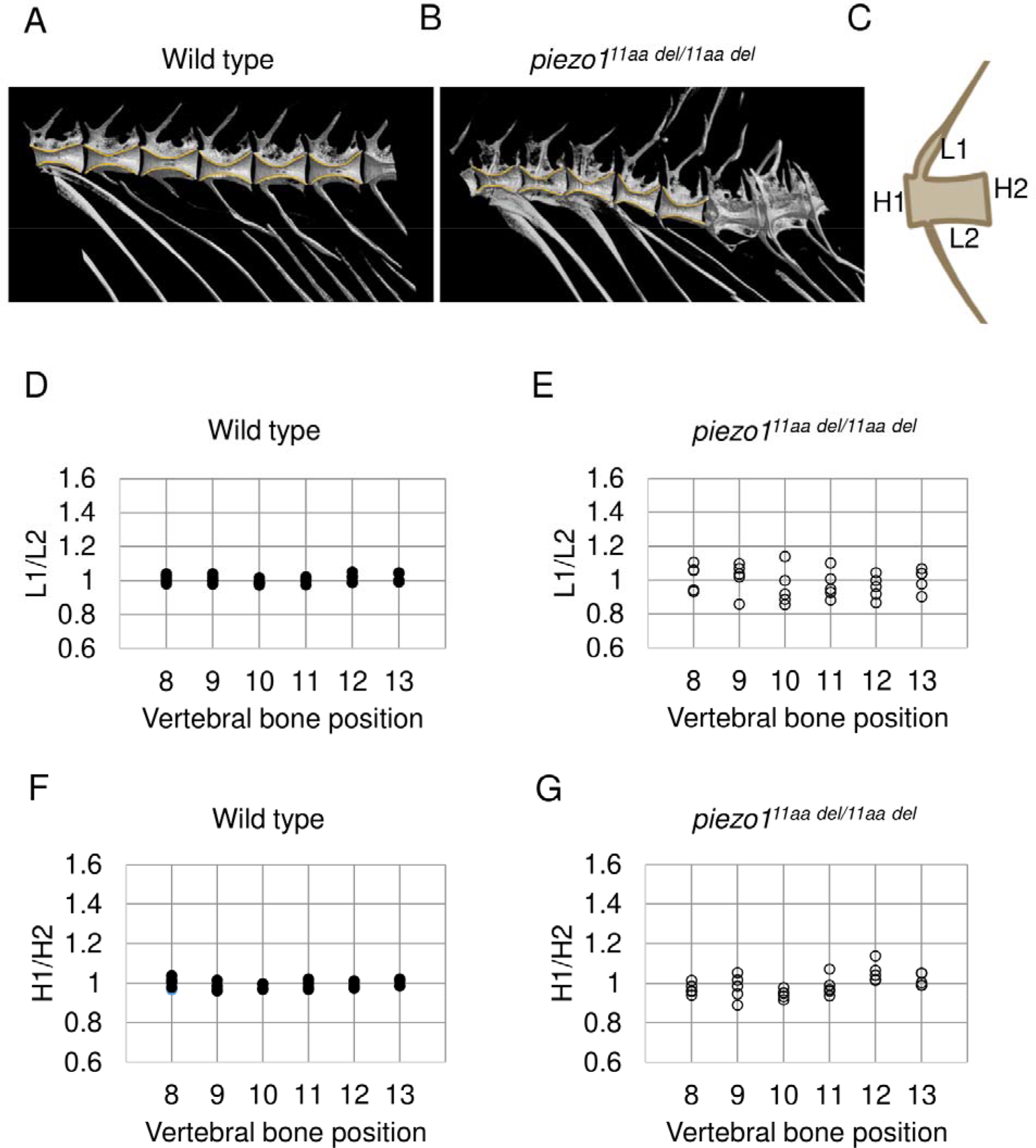
Detailed analysis of vertebra shape. (A)(B) Comparison of 3D reconstruction of micro-CT images of vertebral bone between wildtype and *piezo1*^*11aa del/11aa del*^ at the abdominal part from lateral view. (C) Schematic diagram and quantification of the (D)(E) length ratio (L1/L2) and (F)(G) height ratio (H1/H2) of single vertebrae measurement. About 6 vertebrae from wildtype (N=5) and mutant (N=5) were assessed.

### 3.7 Abnormal intervertebral disc

Several studies have shown that adolescent idiopathic scoliosis can lead to intervertebral disc (IVD) degeneration in later life (Bertram et al., 2006;Hristova et al., 2011;Akazawa et al., 2017). Many of the features described, such as osteophyte formation, endplate sclerosis, facet joint changes, IVD narrowing, and ectopic calcification, have been observed in individuals with intervertebral disc degeneration. In *piezo1*^*11aa del/11aa del*^ mutant fish at the age of 5 months, similar abnormal traits were observed in the intervertebral joint, including IVD calcification (IC), bone fusion (BF), ectopic bone calcification (EC), end-plate sclerosis (ES), and osteophytes (OP) (Figures 5A-5F).

**Figure 5.**
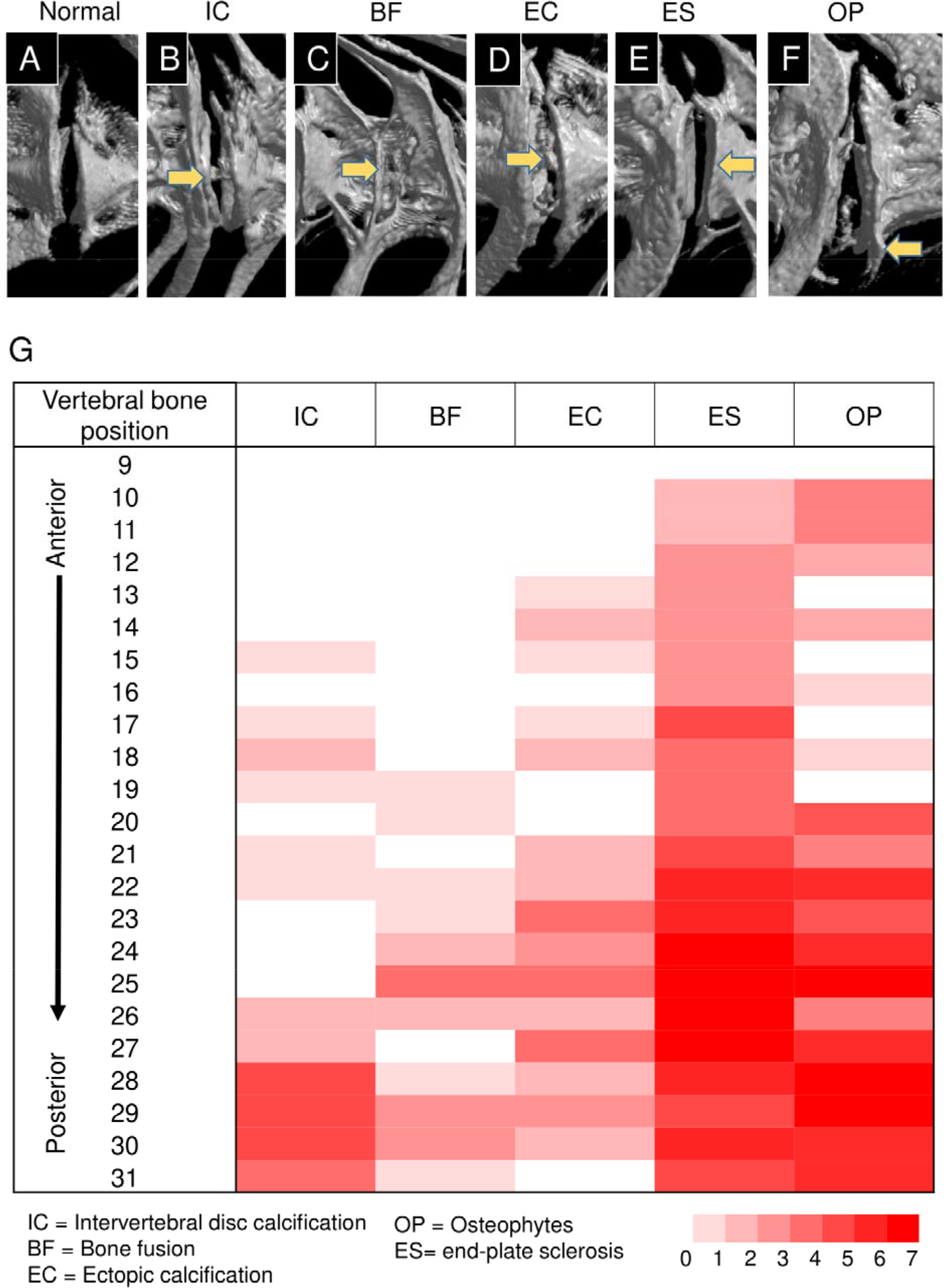
Abnormal changes of vertebral bone and intervertebral disc (IVD) in *piezo1*^*11aa del/11aa del*^. (A) Normal feature of IVD in wildtype; (B) Intervertebral disc calcification (IC); (C) Bone fusion (BF); (D) Ectopic calcification (EC); (E) End-plate sclerosis (ES); (F) Osteophytes formation (OP). (G) Heat map depicting the position and frequency of abnormal features in intervertebral disc region.

As shown in Figure 5G, malformation of vertebrae can occur at any site, regardless of the type of morphological abnormality, but are more frequent at sites close to the tail. Particularly distinct are the intervertebral calcification sites, with more than 80% occurring in the 28th to 30th vertebrae, where the physical stress from tail movement is expected to be greatest. This suggests that the cause of the mutation is mechanical stress.

### 3.8 Introducing functional *piezo1* gene into *piezo1* scoliosis mutant could develop normal spine

Identification of the causative cell is important for use as a disease model. The most likely candidate is osteoblasts, but since the *piezo1* gene is expressed in many tissues, we cannot rule out osteoblasts as the cause of the mutation in *piezo1*^*11aa del/11aa del*^ mutant. Therefore, we attempted to express *piezo1* using the osteoblast-specific promoter *SP7/osterix* to see if the mutation could be rescued (Figure 6A).

**Figure 6.**
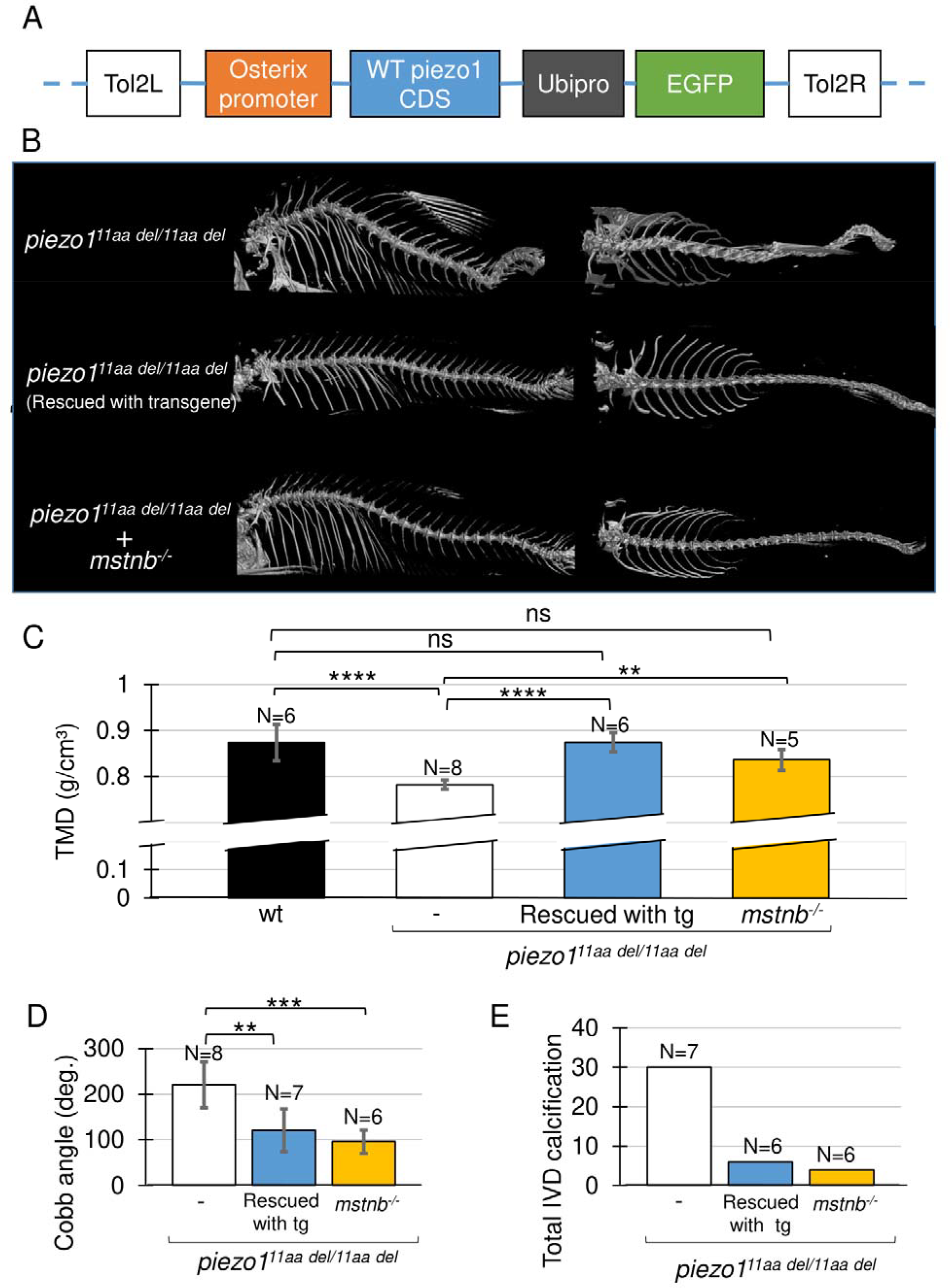
Introducing functional *piezo1* gene or increasing muscle mass can alleviate scoliosis symptoms. (A) Schematic diagram of rescue plasmid. Notably, SP7/osterix promoter was used to express functional *piezo1* gene. In addition, a ubiquitinb promoter (ubipro)-driven EGFP (Enhanced Green Fluorescent Protein) cassette was used to simply the genotyping of fish embryo. (B) Comparison of 3D reconstruction of micro-CT images. (C) Quantification of TMD from 5 individual segments of pre-caudal vertebrae of wildtype (N=6), *piezo1*^*11aa del/11aa del*^ mutant (N=8), rescue mutant fish (N=6), and double mutant of *piezo1*^*11aa del/11aa del*^; *mstnb*^*-/-*^(N=5). (D) Combined cobb angle of *piezo1*^*11aa del/11aa del*^ mutant (N=8) rescue mutant fish (N=7), and double mutant of *piezo1*^*11aa del/11aa del*^; *mstnb*^*-/-*^ (N=6). (E) Graph depicting the total number of intervertebral disc calcifications. Value are presented as mean ± SD and analyzed using one-way ANOVA followed by Tukey’s test.*\*\* P < 0*.*01; *** P < 0*.*001* ; *\*\*\*\* P < 0*.*0001*

Results are shown in Figure 6B, as expected, the mutant fish harboring transgene with the rescue plasmid showed a nearly normal phenotype. Although some small bones were slightly bent along the vertebrae, the reduced phenotype in the rescue mutant was confirmed in all of the TMD, Cobb angle, and total number of IVD calcifications (Figure 6C-6E).

This result confirms that the cells responsible for the bone morphological abnormalities in *piezo1* are osteoblasts. Furthermore, it shows that the severity of the morphological abnormality can be regulated by artificially altering the expression level of *piezo1*. This will increase the utility of the *piezo1*^*11 aa del/11aa del*^ fish as a pathological model.

### 3.9 Increase muscle mass could alleviate scoliosis symptoms

To evaluate the utility of *piezo1*^*11aa del/11aa del*^ mutant fish as a model for pathology, it is necessary to determine whether the abnormal bone morphology of fish responds in the same manner as in humans to manipulations that aggravate or alleviate the symptoms of scoliosis. Scoliosis that develops during adolescence can sometimes be attributed to muscle defects (Lv et al., 2021). Furthermore, exercise, especially weight-bearing exercises that increase muscle mass, has been shown to be an effective treatment for scoliosis (Lau et al., 2021;Hui et al., 2022). Based on these facts, we hypothesized that increasing muscle mass might reduce scoliosis-like symptoms in *piezo1*^*11 aa del/11aa del*^ mutant fish. In fish and mice, inhibition of the *myostatin (mstnb)* gene, a protein that regulates muscle development (Welle et al., 2007;Suh et al., 2020) can increase muscle mass with little effect on other organs. Therefore, disrupting the *myostatin* gene in *piezo1*^*11 aa del/11aa del*^ fish can easily be used to study the relationship between muscle mass and the pathological level of scoliosis.

Using previously published guide RNA (Gao et al., 2016), zebrafish *myostatin* knockouts were produced by CRISPR/Cas9. Zebrafish *mstnb*^*-/-*^ exhibited relatively larger body size compared to wildtype (Gao et al., 2016), with a clear increase in muscle mass around the spine (Supplementary Figure S6). Next, *myostatin* knockout zebrafish were crossed twice with *piezo1*^*11aa del/+*^ to examine the relationship between increased muscle mass and scoliosis severity. Figure 6 shows an overview of the vertebrae and the statistical data. As expected, the strength of the spinal curvature decreased with increasing muscle (Figure 6B), and tissue mineral density increased (Figure 6C). When quantified in terms of Cobb angle, it decreased to almost 30% (Figure 6D). Also, abnormal calcification in the intervertebral disc (IVD) region was also barely observed (Figure 6E; Supplementary Figure S7). These findings are strong evidence that increasing muscle mass alleviates scoliosis and promotes bone formation, suggesting a potential therapeutic approach for the treatment of scoliosis. *piezo1*^*11aa del/11aa del*^ fish may be used effectively as a pathological model for scoliosis. We suggest that *piezo1*^*11aa del/11aa del*^ fish is a good model for the treatment of scoliosis, as they have been shown to be able to increase the muscle mass of the spine and to increase the bone mineral density of the spine.

## 4. Discussion

Scoliosis is a disease caused by the almost unique posture of humans in the vertebrate world, in which they stand upright on two legs, and it has been difficult to obtain an experimental system that can serve as a model for the disease (Castelein et al., 2005;Bobyn et al., 2015;Xie et al., 2022). On the other hand, there have been cases of scoliosis-like symptoms in fish (Gorman et al., 2007;Gorman and Breden, 2009), and it is thought that the longitudinal compression of the spine caused by the propulsive force of the caudal fin is similar to that caused by the upright posture in humans. We have established a pathological model of scoliosis in zebrafish by manipulating the *piezo* genes, which are mechanotropic stimulus receptors in the cells. Zebrafish have two *piezo* genes (*piezo1* and *piezo2a*) involved in bone formation. When both genes were knocked out, various abnormalities including bone formation appeared in early development and early death occurred. However, an in-frame mutation in the *piezo1* gene (11 amino acid deletion) was not lethal and caused morphological abnormalities of the spine similar to scoliosis in humans. The fact that the morphological abnormalities occur later in life and that bone density is reduced, also suggests that the zebrafish spinal morphology may be homologous to human scoliosis. Since zebrafish can be easily bred in large numbers and genetic manipulation tools have been developed, it is expected that mutant zebrafish will be used to study the pathogenesis of scoliosis and to screen for therapeutic agents.

### 4.1 Usefulness as a pathological model

The pathological model of scoliosis created in this study using zebrafish has several advantages besides the ease of rearing a large number of animals. As shown in Figure 6, the severity of the disease can be modulated by the expression of normal *piezo1* and *piezo2a* genes. This could be a useful property when screening for therapies. When screening for therapeutic agents, it is expected that if the severity of the model is too strong, the effect will not be apparent, and if the severity is too weak, the data may be unstable. Screening for individuals of varying severity would reduce the likelihood of missing an effective drug. Another advantage is the ability to test the efficacy of treatments other than relaxing agents, such as physical therapy, as in the present study, where muscle augmentation due to deletion of the *myostatin* gene induced a resolution of symptoms. A more advanced approach would be to investigate the relationship between exercise therapy and scoliosis, such as artificially inducing muscle contraction with light.

### 4.2 Causal relationship between scoliosis and osteoporosis

Although scoliosis and osteoporosis are distinct skeletal conditions with separate sets of symptoms and underlying causes, there is a suggested connection between them. Numerous clinical reports have indicated that individuals with idiopathic scoliosis often exhibit lower bone mineral density (BMD) (Li et al., 2008;Sadat-Ali et al., 2008;Nishida et al., 2023;Wu et al., 2023), a key indicator of osteoporosis. Similar to these studies, we also reported remarkably lower BMD values in *piezo1*^*11aa del/11aa del*^ mutant compared to siblings. However, it remains unclear whether osteoporosis plays a causative role or is a consequence of scoliosis.

Based on current knowledge, it is plausible that osteoporosis may be an underlying factor contributing to the development of scoliosis. Histological examinations of trabecular bone in idiopathic scoliosis patients have shown decreased osteoblast activity (Cheng et al., 2001). Additionally, children diagnosed with idiopathic scoliosis have demonstrated diminished osteogenic differentiation potential in their mesenchymal stem cells (Park et al., 2009;Chen et al., 2016). These findings suggest that the decline in osteoblast activity, leading to a reduction in BMD, may occur prior to the formation of lateral spinal curvature. Low bone density resulting from abnormal osteogenic activity could increase the risk of bone fractures. The presence of microfractures may, in turn, contribute to bone asymmetry, potentially exacerbated by axial loading. Eventually, this leads to the development of spinal deformity in a cyclical and interdependent manner.

### 4.3 *piezo1*^*11aa del/11aa del*^ mutant for a tool to investigate the *piezo* genes in other cells

*piezo1* and *piezo2a* are homologous in function and both genes are expressed in almost all cells. Double deletion mutations cause hypoplasia in bladder and body segments as well as osteogenesis imperfecta. These results indicate that the *piezo* genes are also essential for the formation of other organs in zebrafish and are consistent with other reports showing *piezo* gene function in a variety of cells. Although 11aa del-Piezo1 is a mutation in the *piezo1* gene, it also reduces the function of *piezo2a* with a dominant-negative effect. This feature may allow in vivo studies of *piezo* genes’ function in other cells. Specifically, expression of the 11aa del-Piezo1 mutant gene using a target cell-specific promoter would avoid lethality and reveal the role of the *piezo* genes in those cells.

## Conflict of Interest

The authors declare that the research was conducted in the absence of any commercial or financial relationships that could be construed as a potential conflict of interest.

## Author Contributions

R: Conceptualization, Investigation, Data curation, Methodology, Visualization, Writing – original draft. TA: Conceptualization, Methodology, Supervision, Validation, Writing – review and editing. MW: Conceptualization, Methodology, Validation, Writing – review and editing. SK: Conceptualization, Funding acquisition, Supervision, Resources, Investigation, Writing – review and editing.

## Funding

This research was supported by Grant-in-Aid for Transformative Research Areas (20H05943) from the Ministry of Education, Culture, Sports, Science and Technology (MEXT), Japan, and Grant-in-Aid for Scientific Research (19H00994) from Japan Society for the Promotion of Science (JSPS).

## Acknowledgments

The authors would like to thank the members of the Kondo laboratory for their experimental support and for providing valuable feedback on this study. Piezo1-deficient N2A cell was gifted from Dr. Kenta Maruyama at National Institute for Physiological Sciences & Dr. Yasunori Takayama at Showa University.

## Notes

### Competing Interest Statement

The authors have declared no competing interest.

