## supplementary table S1 for "Piezo1 Mutant Zebrafish as a Model of Idiopathic Scoliosis"

### 1 Supplementary Tables

**Table S1**: Oligo sequences used for CRISPR-Cas9 gRNA generation

| Gene | Primers name | Sequences |
|---|---|---|
| *piezo1* | piezo1 sgRNA exon 5 | GCT AAT ACG ACT CAC TAT AGG GAG CCA CAG CAC ACC CTG GTT TTA GAG CT |
| *piezo2a* | piezo2a sgRNA Exon 4 | GCT AAT ACG ACT CAC TAT AGG GCC ACG CTC ATC CGC CTC GTT TTA GAG CT |
| *piezo1* in-frame mutant (11aa del) | piezo1-sg1F | TAG GAG CGA AAT ATG CAG GCT G |
| | piezo1-sg1R | AAA CCA GCC TGC ATA TTT GCG T |
| *mstnb* | mstnb-sg1-F | TAG GAG CCT TCC ACA GCC ACG G |
| | mstnb-sg1-R | AAA CCC GTG GCT GTG GAA GGC T |

**Table S2**: Primers used for genotyping

| Gene | Primers | Sequences |
|---|---|---|
| *piezo1* | piezo1 int-4F | TCC TGG GAC GTA ACA AAG CA |
| | piezo1 int-5R | AGG CCC AGA CTA ACA GCA TT |
| *piezo2a* | piezo2a int-4F | TTT GAC AAC AAA ATG ACT CAC TAA TTG |
| | piezo2a int-5R | CGA TGT ACA AAA AGC CAC CA |
| *piezo1* in-frame mutant (11aa del) | piezo1 int-1F | AAA ATC ACA GCA GGG TGA AT |
| | piezo1 int-2R | GGC AGA CTA TTG CAA CAT TGA |
| *mstnb* | mstnb-ex1F | ACA TCC TTT AGC ACG CCT TG |
| | mstnb-int1R | CTG CGT AAA GGG TCT CTC CA |

**Table S3**: Primers used for qPCR

| Gene | Primers | Sequences |
|---|---|---|
| *piezo1* | piezo1 QPF | GAG AGG ATG CGG CTT CTC AA |
| | piezo1 QPR | CCA CAT GGT GAA TCC GTC CA |

### 2 Supplementary Figures

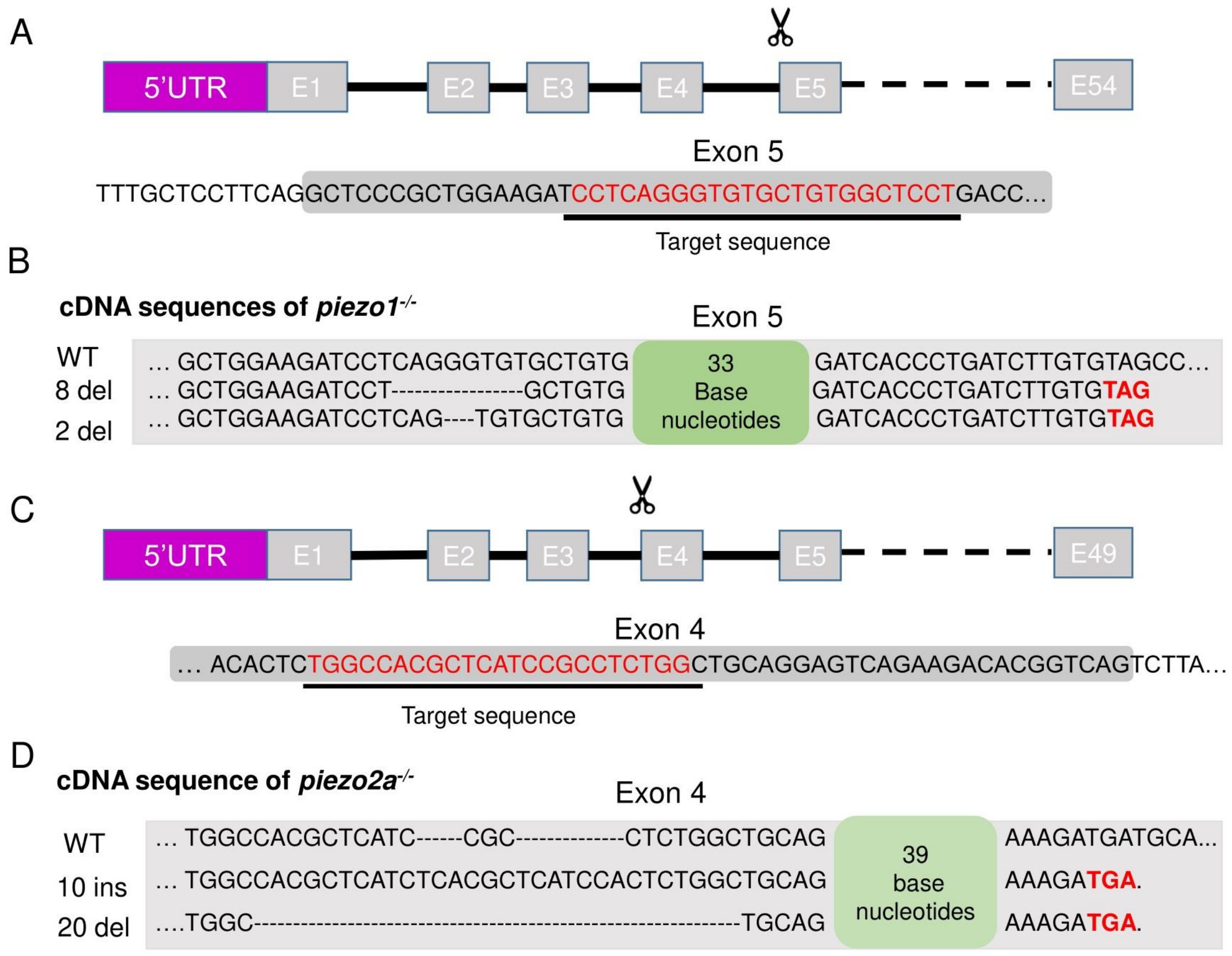


**Supplementary Figure S1.** Generation of *piezo1*$^{-/-}$ and *piezo2a*$^{-/-}$**.** (A)(C) Schematic diagram of CRISPR/Cas9 targeting of *piezo1* and *piezo2a* genes by guide RNAs targeting the N-terminal regions of *piezo1* exon 5 and *piezo2a* exon 4, respectively. (B)(D) sequence confirmation of *piezo1*$^{-/-}$ alleles and *piezo2a*$^{-/-}$ alleles showing premature terminator codon.

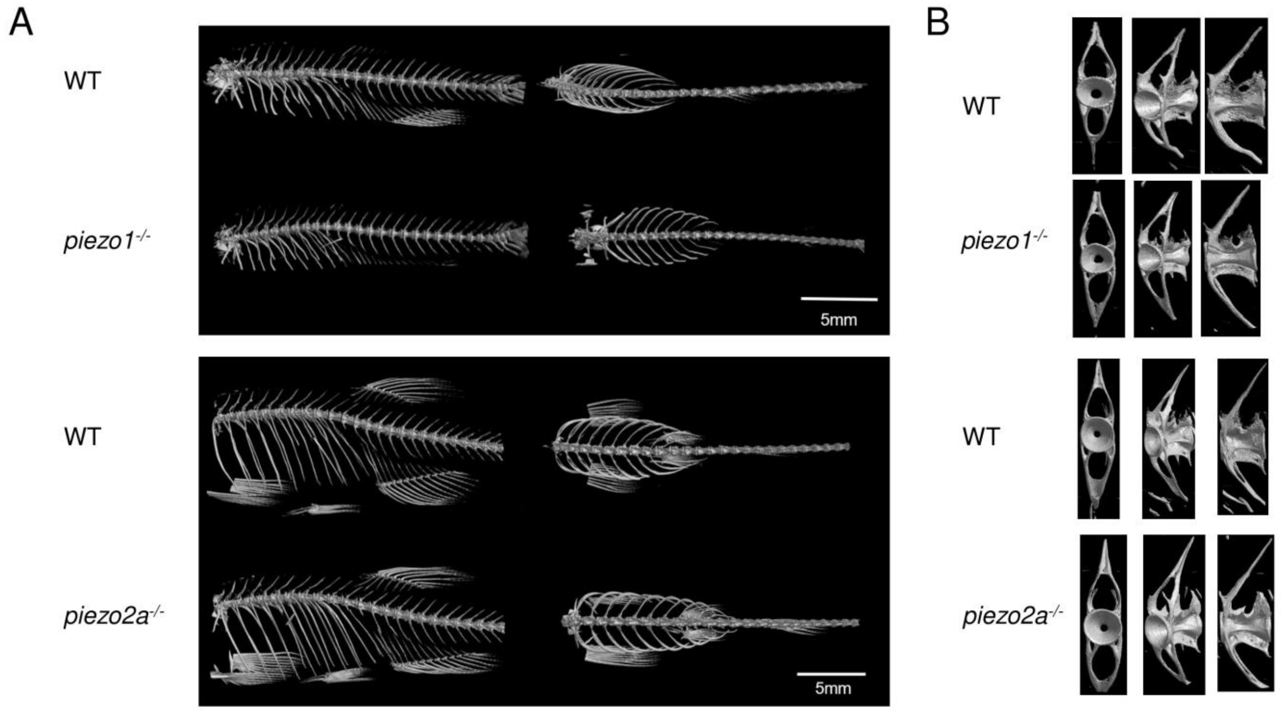


**Supplementary Figure S2.** Bone phenotypes of *piezo1*$^{-/-}$ and *piezo2a*$^{-/-}$. Comparison of 3D reconstruction of micro-CT images of (A) whole body and (B) bone segment between wildtype and *piezo1*$^{-/-}$ at 4 mpf (N=6) also wildtype and *piezo2a*$^{-/-}$ at 6 mpf (N=4). Notably, there was no morphological abnormalities in each null mutant.

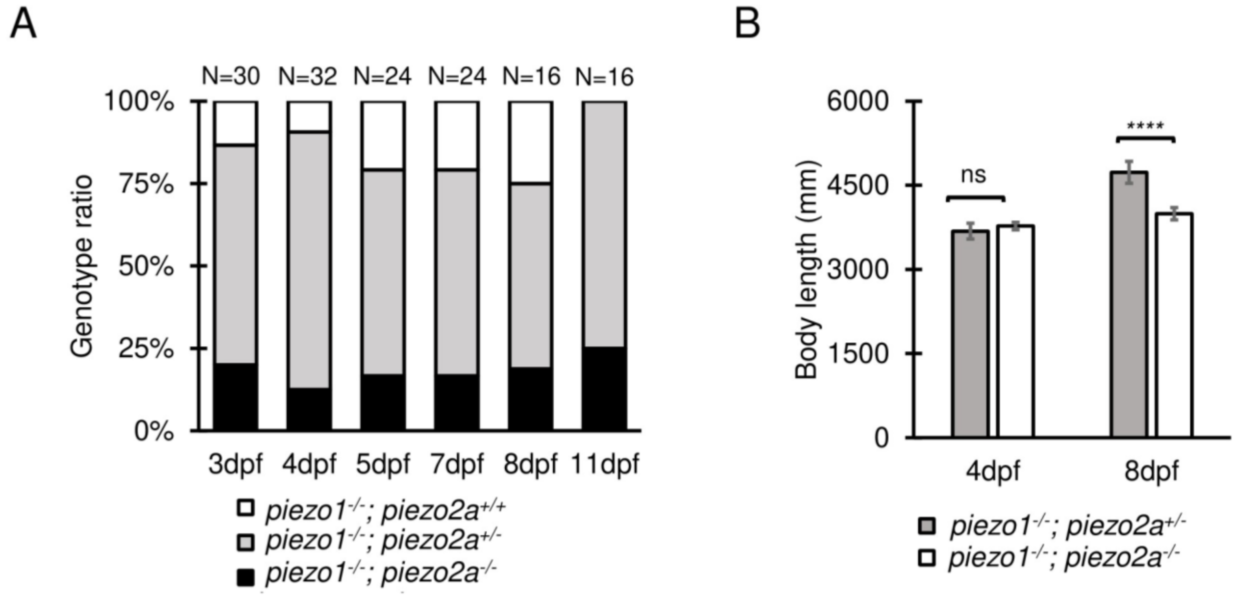


**Supplementary Figure S3.** Survival rate and total body length of *piezo1*$^{-/-}$; *piezo2a*$^{-/-}$. (A) Genotype ratio between double knock out and sibling in several time points. (B) Graph depicting total body length at 4dpf and 8dpf (N=8 for each group). Values are presented as mean ± SD and analyzed using student t-test. **** *P < 0.0001*. ns indicates not significant.

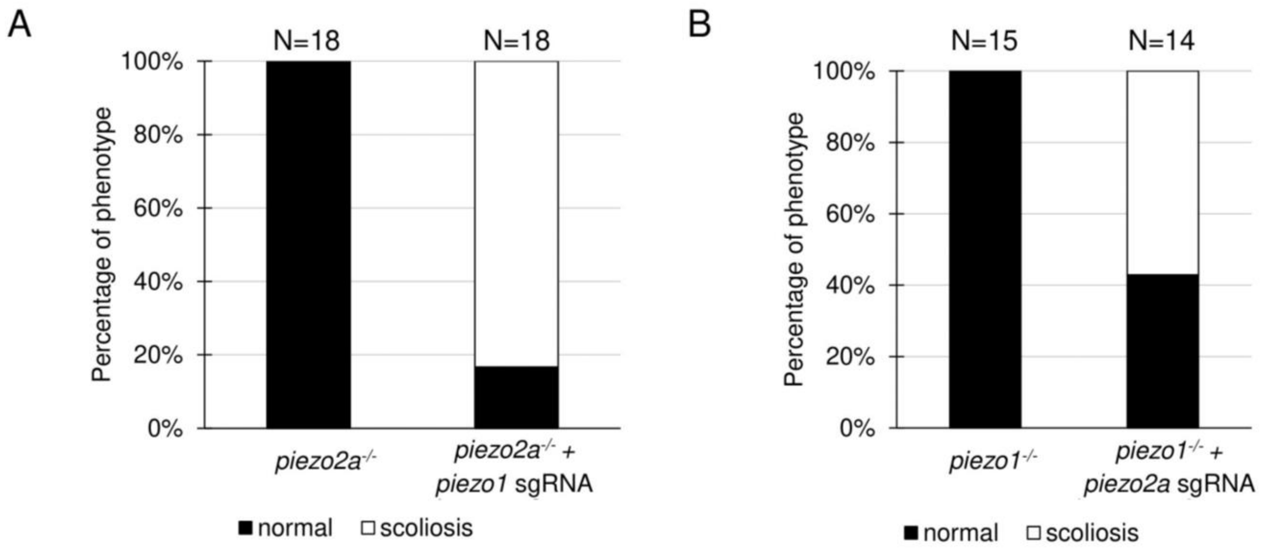


**Supplementary Figure S4.** Percentages of scoliosis phenotype in mosaic mutants.

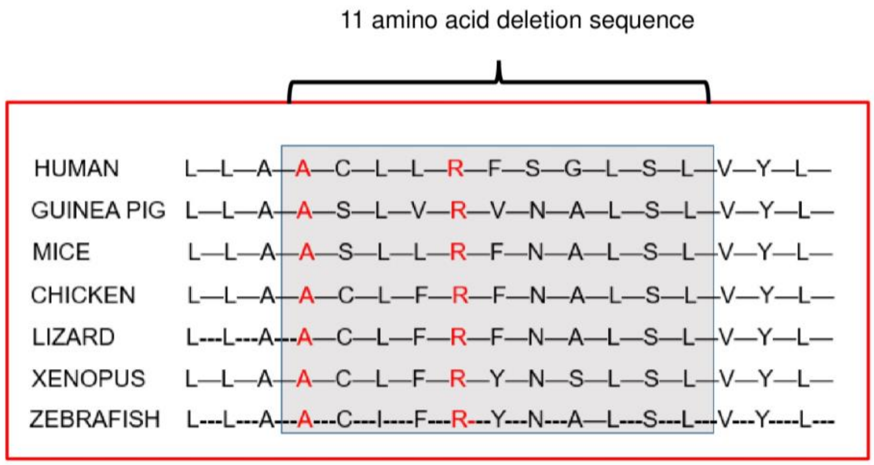


**Supplementary Figure S5.** 11 amino acid deletion region in Piezo1 is highly conserved among vertebrates.

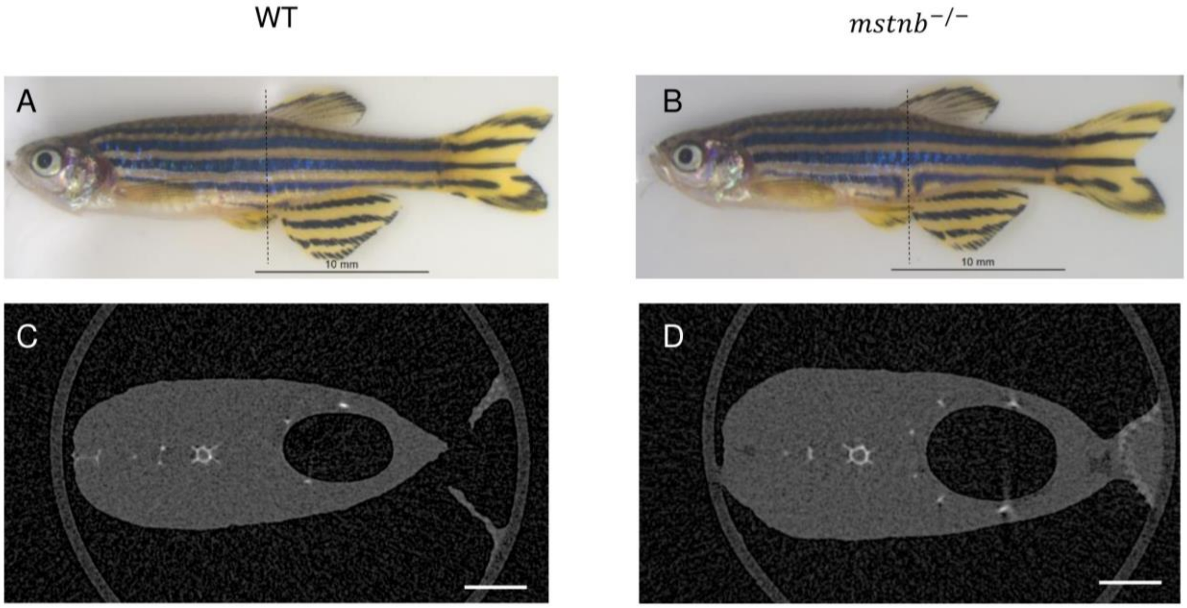


**Supplementary Figure S6.** Phenotype of *mstnb*$^{-/-}$. (A)(B) Gross phenotype between wildtype and *mstnb*$^{-/-}$ at 4 months after fertilization. Notably, the body size of *mstnb*$^{-/-}$ was slightly larger. (C)(D) Cross-sections view of abdominal part. Used scale is 50 μm.

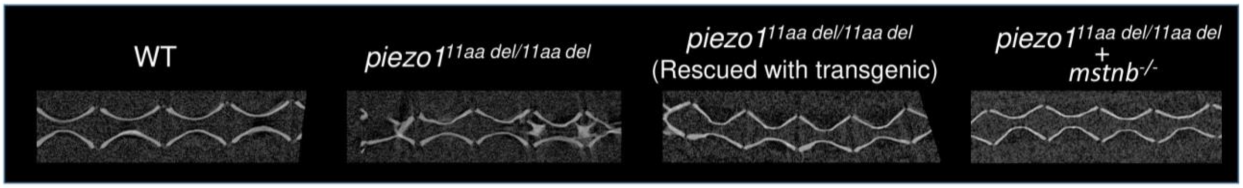


**Supplementary Figure S7.** Sagittal section of the spine. Rescue mutant and *piezo1*$^{11aa\ del/11aa\ del}$*; mstnb*$^{-/-}$ have reduced intervertebral disc calcification.
